# A mouse-adapted SARS-CoV-2 model for the evaluation of COVID-19 medical countermeasures

**DOI:** 10.1101/2020.05.06.081497

**Authors:** Kenneth H. Dinnon, Sarah R. Leist, Alexandra Schäfer, Caitlin E. Edwards, David R. Martinez, Stephanie A. Montgomery, Ande West, Boyd L. Yount, Yixuan J. Hou, Lily E. Adams, Kendra L. Gully, Ariane J. Brown, Emily Huang, Matthew D. Bryant, Ingrid C. Choong, Jeffrey S. Glenn, Lisa E. Gralinski, Timothy P. Sheahan, Ralph S. Baric

## Abstract

Coronaviruses are prone to emergence into new host species most recently evidenced by SARS-CoV-2, the causative agent of the COVID-19 pandemic. Small animal models that recapitulate SARS-CoV-2 disease are desperately needed to rapidly evaluate medical countermeasures (MCMs). SARS-CoV-2 cannot infect wildtype laboratory mice due to inefficient interactions between the viral spike (S) protein and the murine ortholog of the human receptor, ACE2. We used reverse genetics to remodel the S and mACE2 binding interface resulting in a recombinant virus (SARS-CoV-2 MA) that could utilize mACE2 for entry. SARS-CoV-2 MA replicated in both the upper and lower airways of both young adult and aged BALB/c mice. Importantly, disease was more severe in aged mice, and showed more clinically relevant phenotypes than those seen in hACE2 transgenic mice. We then demonstrated the utility of this model through vaccine challenge studies in immune competent mice with native expression of mACE2. Lastly, we show that clinical candidate interferon (IFN) lambda-1a can potently inhibit SARS-CoV-2 replication in primary human airway epithelial cells *in vitro*, and both prophylactic and therapeutic administration diminished replication in mice. Our mouse-adapted SARS-CoV-2 model demonstrates age-related disease pathogenesis and supports the clinical use of IFN lambda-1a treatment in human COVID-19 infections.

---

Zoonotic coronaviruses readily traffic into new host species, as evidenced by the emergence of SARS-CoV in 2002-2003, Middle East Respiratory Coronavirus in 2012 and SARS-CoV-2, the causative agent of the COVID-19 pandemic in 2019. COVID-19 infections have caused millions of infections and hundreds of thousands of deaths worldwide. As there are currently no approved vaccines and only one antiviral approved for emergency use for SARS-CoV-2^1^, small animal model systems are vital to better understand COVID-19 disease mechanisms and to evaluate medical counter measures (MCMs) for improved global health. Mouse models not only provide key insights into the pathogenic mechanisms of CoV disease but can serve as high-throughput preclinical evaluation platforms to identify high performance antivirals and vaccines essential for downstream human trials^2,3^. SARS-CoV-2 enters host cells through the binding of the cellular receptor angiotensin-converting enzyme 2 (ACE2). Unfortunately, standard laboratory mice do not support infection with SARS-CoV-2 due to incompatibility of the S protein to the murine ortholog (mACE2) of the human receptor, complicating model development^4^.

## Remodeling the SARS-CoV-2 mACE2 binding interface

Animal models are critical for development of MCMs to combat the COVID-19 pandemic. While laboratory mice infected with mouse adapted strains of SARS-CoV-1 and MERS CoV have informed our understanding of viral pathogenesis and intervention strategies, the murine ortholog of the human SARS-CoV-2 receptor, mACE2, does not facilitate infection. To determine the utility of *hACE2* transgenic mice as a model for SARS-CoV-2 disease, we infected epithelial cell-specific *HFH4* promoter driven *hACE2*-overexpressing mice with SARS-CoV-2^5^. *hACE2* mice lost minimal weight yet only 60% survived by 5 days post infection (dpi) (Fig. S1A-B). Virus replication was observed in the lung but not in the (Fig. S1C). Similar to previously reported SARS-CoV-1 infection of *hACE2* mice, SARS-CoV-2 replication was detected in the brains of those that succumbed to infection suggesting that mortality was driven by viral neuroinvasion (Fig. S1 D). Lastly, whole body plethysmography (WBP) reveals pulmonary function remained at normal levels for the duration of these studies proving further evidence that respiratory infection was likely not a major driver of mortality (Fig. S1F-G). Thus, in the presence of a *hACE2* transgene, mice fully support SARS-CoV-2 replication but the observed pathogenesis failed to accurately model the disease course seen in humans.

Rather than alter the host, we next sought to remodel the SARS-CoV-2 spike receptor binding domain to facilitate efficient binding to mACE2 and therefore productive viral infection. Upon comparing the ACE2 contact residues in the RBDs of several group 2B coronaviruses capable of infecting mice (i.e. SARS-CoV-1, WIV1, and SHC014) to those of SARS-CoV-2, residue 498 of SARS-CoV-2 was uniquely divergent (Fig. 1A, Fig. S2). In addition, molecular modeling of the SARS-CoV-2 RBD and receptor interface revealed a loss of interaction between Q498 of the SARS-CoV-2 spike and Q42 of mACE2 (Fig 1B-C), which may diminish binding efficiency. Thus, we predicted the substitution of residue Q498, and adjacent P499, with those from WIV1 and SARS-CoV-1 would restore the interaction with Q42 of mACE2, while preserving interaction with hACE2 (Fig. 1D-E). Using reverse genetics, we engineered Q498T/P499Y into the SARS-CoV-2 S gene and recovered the recombinant virus (SARS-CoV-2 MA). Importantly, SARS-CoV-2 MA replicated to similar levels as the parental WT virus in Vero E6 cells (Fig. 1F) and unlike WT virus, SARS-CoV-2 MA could infect cells expressing mACE2 (Fig. 1G).

**Figure 1.**
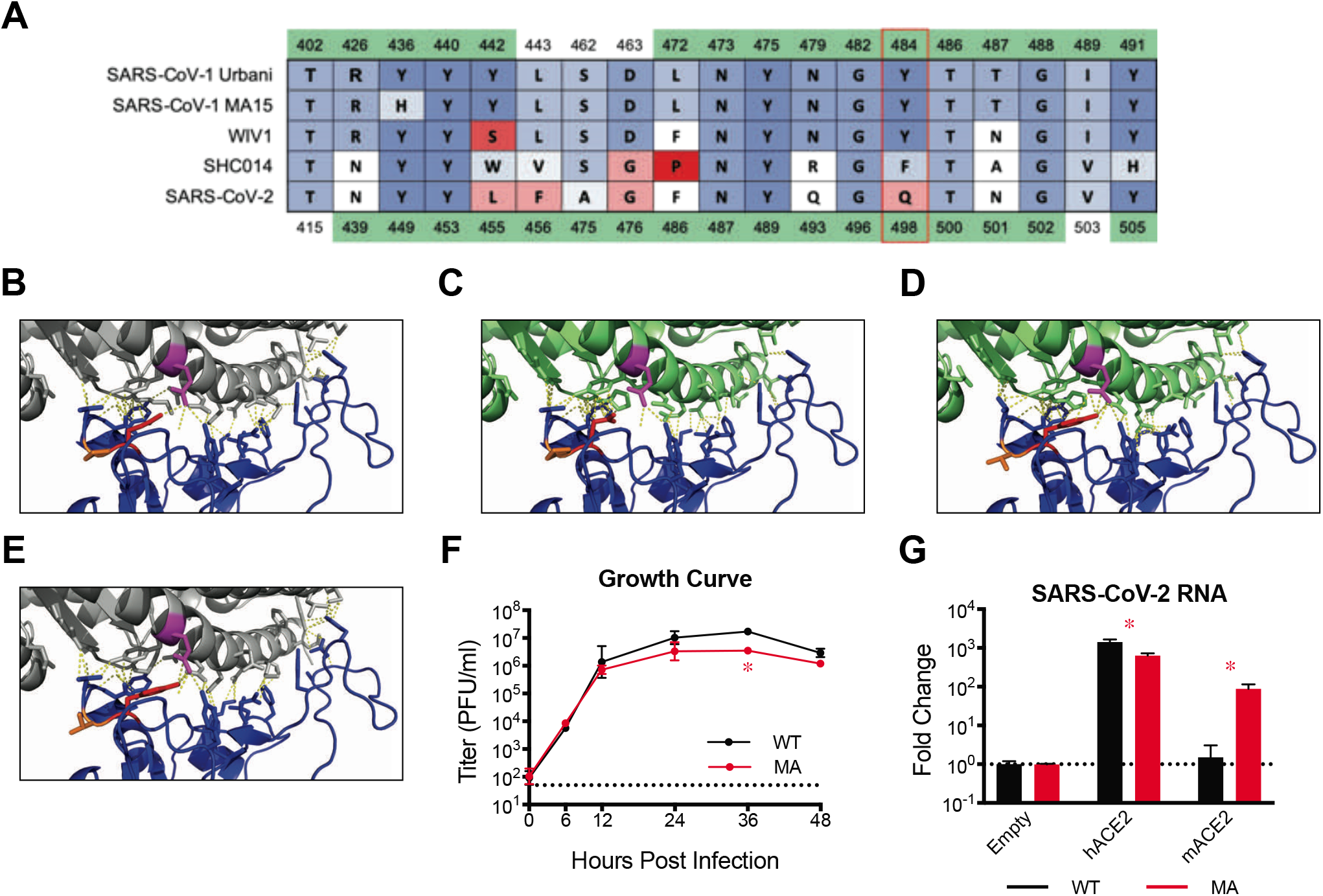
Generation of mouse adapted SARS-CoV-2 MA. **(A)** Amino acid table of group 2B spike receptor binding domains (RBDs). Amino acid positions are numbered above in reference to SARS-CoV-1, and below in reference to SARS-CoV-2. Green highlighted residues are hACE2 contacts as determined by published crystal structures. Amino acids colored by BLOSUM62 conservation score relative to SARS-CoV-1 Urbani, with red being least conserved to blue being most conserved. SARS-CoV-1 Urbani, SARS-CoV-1 MA15, WIV1, and SHC014 can utilize mACE2 as a functional receptor whereas SARS-CoV-2 cannot. Red box indicates residue Q498 in SARS-CoV-2 that is a hACE2 contact for both SARS-CoV-1 and SARS-CoV-2, and is uniquely divergent in SARS-CoV-2. **(B)** SARS-CoV-2 WT RBD (blue ribbon structure) and hACE2 (gray ribbon structure) interface (PDB: 6m0j). SARS-CoV-2 Q498 (red) interacts with Q42 (magenta) of hACE2. **(C)** Modelling of SARS-CoV-2 WT RBD (blue ribbon structure) and mACE2 (green ribbon structure). SARS-CoV-2 Q498 (red) no longer interacts with Q42 (magenta) of mACE2. **(D)** Modelling of SARS-CoV-2 Q498T (red) and P499Y (orange) substitutions restore interaction with Q42 of mACE2. **(E)** Modelling of SARS-CoV-2 Q498T/P499Y maintains interaction with Q42 of hACE2. **(F)** Single step growth curve of SARS-CoV-2 WT and SARS-CoV-2 MA in Vero E6 cells (n=3 for each group, serially sampled). Dotted line represents limit of detection. Log transformed data analyzed by 2-factor ANOVA followed by Sidak’s multiple comparisons. **(G)** Non-permissive DBT-9 cells were transfected to express hACE2 or mACE2 and infected with SARS-CoV-2 WT and SARS-CoV-2 MA. Viral RNA was quantified by qRT-PCR and scaled to empty vector transfected cells (n=3 for each group). Dotted line represents fold change of 1. Log transformed data analyzed by 2-factor ANOVA followed by Dunnett’s multiple comparisons. The line represents the mean and error bars represent standard deviation. Asterisk denotes p<0.05.

## Mouse adapted SARS-CoV-2 replicates in the upper and lower airways of WT mice

After demonstrating SARS-CoV-2 MA could utilize mACE2 for entry, we sought to determine if this virus could infect young adult WT mice. While overt clinical signs of infection (i.e. weight loss) were not observed in young adult BALB/c mice infected with 10^5^ PFU SARS-CoV-2 MA (Fig. 2A), high titer virus replication (6.93×10^5^ PFU/tissue) was noted in lung tissue on 2dpi but was cleared by 4dpi (Fig. 2B). Under identical conditions, SARS-CoV-2 did not replicate in mice. Histological analysis of SARS-CoV-2 MA infected mice revealed inflammation of small conducting airways on 2dpi, associated with high levels of viral antigen staining (Fig. S3A). Viral replication was limited to conducting airways and absent in interstitium, and is absent on 4dpi, concordant with clearance of viral titer (Fig. 2B, Fig. S3B). Similar to that observed in humans, SARS-CoV-2 MA replication was also observed in the upper airway although the magnitude was less than that seen in the lung (Fig. 2C). The loss of pulmonary function as measured by WBP is commonly observed in murine models of emerging CoV pathogenesis^6,7^ and is an important clinically relevant measure of disease. Using WBP, we evaluated several complementary metrics of pulmonary obstruction and bronchoconstriction including PenH and Rpef. Interestingly, SARS-CoV-2 MA young infected mice showed a small but significant change in PenH (Fig. 2D) and a significant decrease in Rpef (Fig. 2E) on 2dpi indicative of a decrease in lung function. Thus, like often seen in young adult humans, infection of young adult mice with SARS-CoV-2 MA resulted in efficient virus replication in the upper and lower airways, limited replication in the parenchyma and was associated with mild to moderate disease.

**Figure 2:**
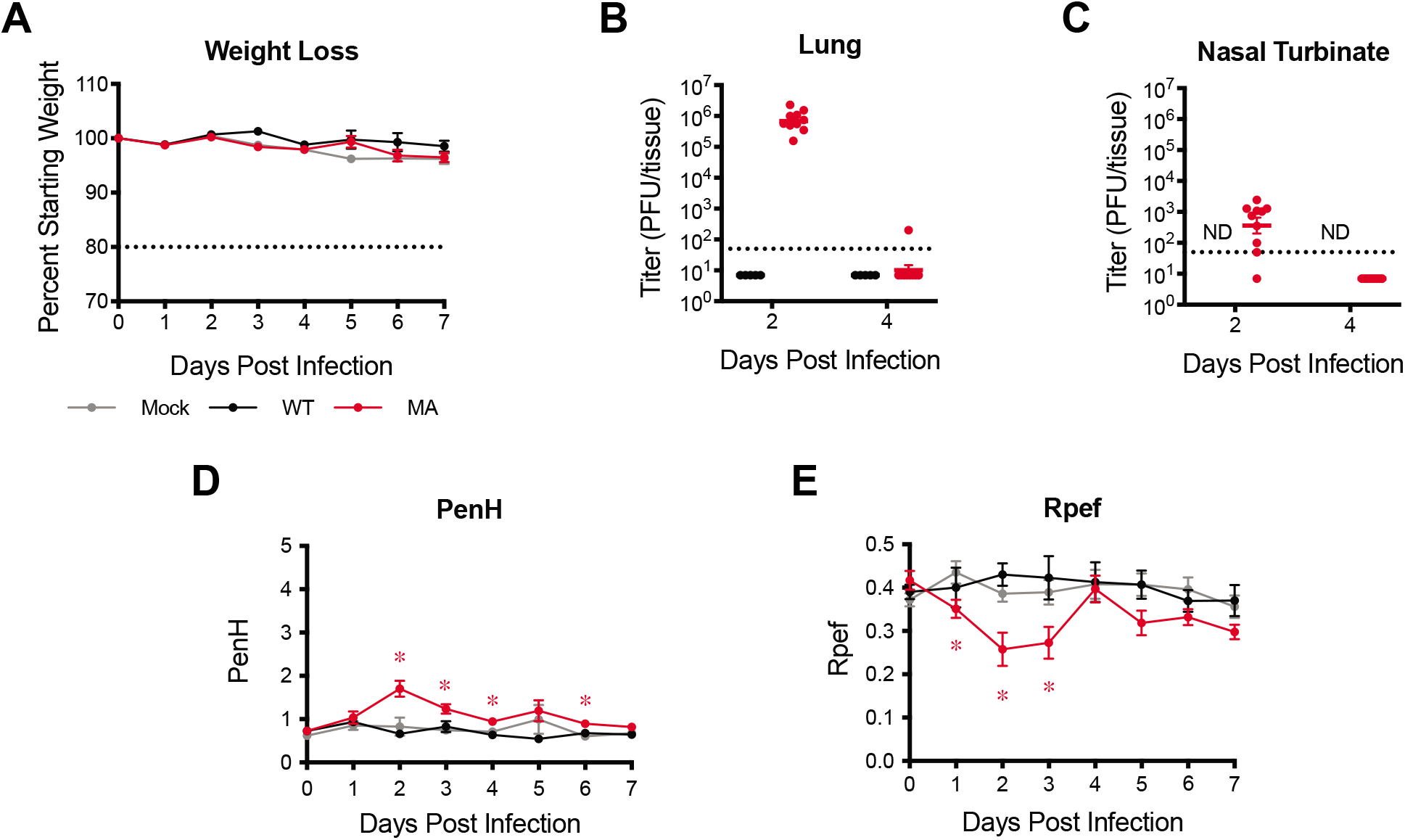
SARS-CoV-2 MA replicates in young BALB/c mice. 12-week-old female BALB/c mice were mock infected (gray), or infected with 10^5^ PFU SARS-CoV-2 WT (black) or MA (red). n=27, 15, 33, respectively. Mice were harvested on day 2, 4 and 7 after infection (n=5-14 per time point). Data combined from two independent experiments. **(A)** Percent starting weight. Dotted line represents weight loss criteria for humane euthanasia. Data analyzed by mixed effects analysis followed by Dunnett’s multiple comparisons. **(B)** Viral lung titer. Dotted line represents limit of detection. Undetected samples are plotted at half the limit of detection. Log transformed data analyzed by 2-factor ANOVA followed by Sidak’s multiple comparisons. **(C)** Nasal turbinate titer. Dotted line represents limit of detection. ‘ND’ denotes titers not determined. Undetected samples are plotted at half the limit of detection. **(D-E)** Whole body plethysmography assessing pulmonary function for PenH **(D)** and Rpef **(E)**. Data analyzed by 2-factor ANOVA followed by Dunnett’s multiple comparisons. The line represents the mean and error bars represent standard error of the mean. Asterisk denotes p<0.05.

## Modeling age-related exacerbation of SARS-CoV-2 pathogenesis in mice

Higher morbidity and mortality rates have been consistently observed in older human populations throughout the COVID-19 pandemic^8^. Additionally, wildtype and mouse adapted SARS-CoV-1 shows strong age dependent disease phenotypes in humans and mice, respectively^9,10^. To determine if the age-related increase in pathogenesis observed in SARS-CoV-2 infected humans would translate to infection of aged mice, we infected 12-month-old BALB/c mice with SARS-CoV-2 MA. In contrast to young adult mice, aged BALB/c mice exhibited a transient yet significant decrease in body weight by 3dpi, which was recovered by 4dpi (Fig. 3A). Old mice also had high titers at 2dpi (1.07×10^6^ PFU/tissue) and detectable virus titers at 4dpi. Similarly, replication in the upper airway persisted in half of the mice at 4dpi (Fig. 3C). Compared to young mice, SARS-CoV-2 MA infected old mice displayed increased inflammation in the lung at 2dpi and 4dpi, and viral antigen was found in both conducting airway epithelium and interstitium (Fig S4A-B). Additionally, at 4dpi, SARS-CoV-2 MA infection induced hemorrhage and formation of bronchus-associated lymphoid tissue (BALT) (Fig. S4B). Lastly, the loss of pulmonary function was more pronounced in aged animals as evidenced by significant differences in PenH and Rpef among mock and SARS-CoV-2 MA infected animals (Fig. 3D-E).

**Figure 3.**
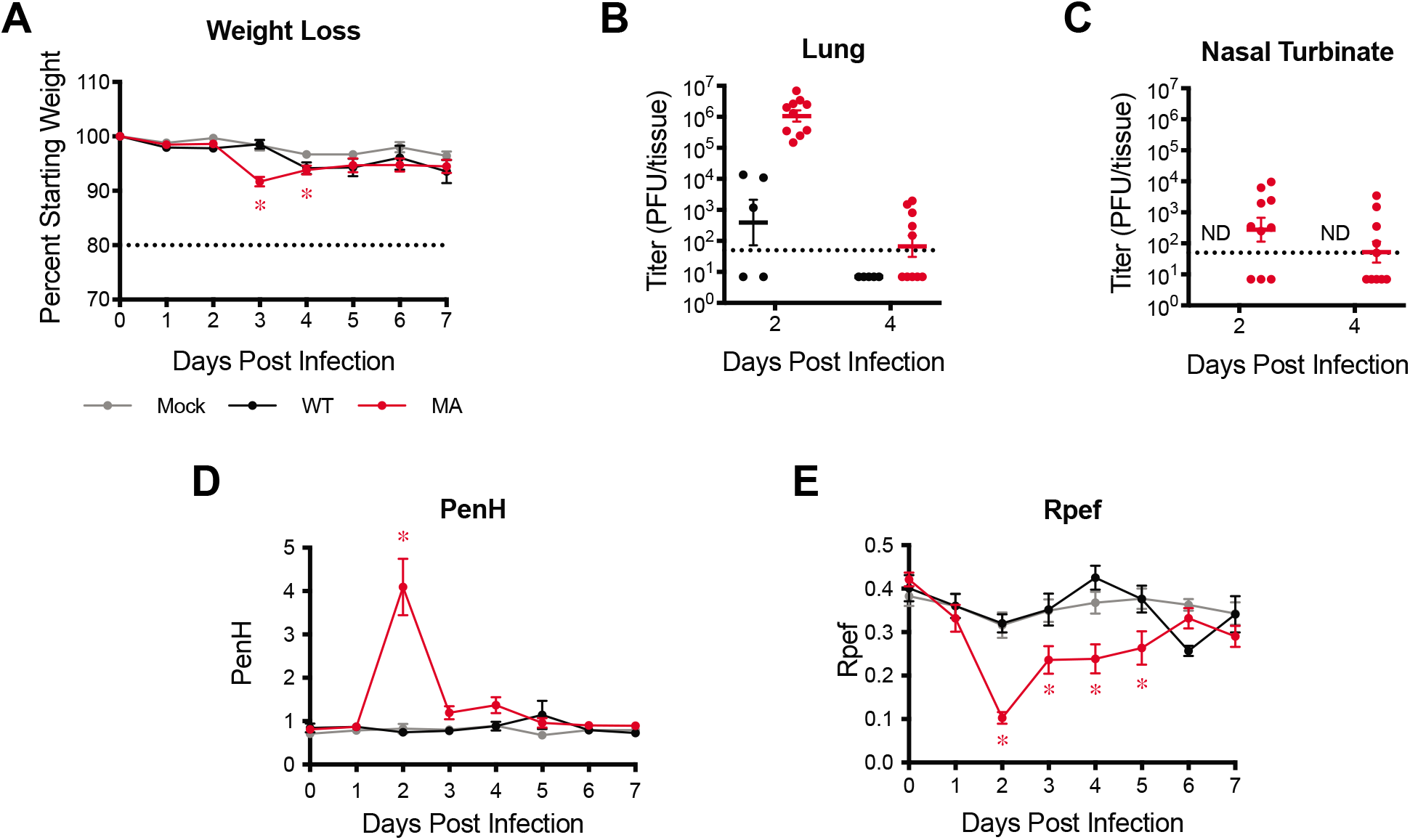
SARS-CoV-2 MA replicates in old BALB/c mice with minor disease. 1-year-old female BALB/c were mock infected (gray), or infected with 10^5^ PFU SARS-CoV-2 WT (black) or MA (red). n=25, 15, 34, respectively. Mice were harvested on day 2, 4 and 7 after infection (n=5-14 per time point). Data combined from two independent experiments. **(A)** Percent starting weight. Dotted line represents weight loss criteria for humane euthanasia. Data analyzed by mixed effects analysis followed by Dunnett’s multiple comparisons. **(B)** Viral lung titer. Dotted line represents limit of detection. Undetected samples are plotted at half the limit of detection. Log transformed data analyzed by 2-factor ANOVA followed by Sidak’s multiple comparisons. **(C)** Nasal turbinate titer. Dotted line represents limit of detection. ‘ND’ denotes titers not determined. Undetected samples are plotted at half the limit of detection. **(D-E)** Whole body plethysmography assessing pulmonary function for PenH **(D)** and Rpef **(E)**. Data analyzed by 2-factor ANOVA followed by Dunnett’s multiple comparisons. The line represents the mean and error bars represent standard error of the mean. Asterisk denotes p<0.05.

## Vectored vaccine and Interferon Lambda-1a therapeutic efficacy against SARS-CoV-2

As demonstrated with mouse adapted strains of SARS-CoV-1^11-15^, a replication competent SARS-CoV-2 MA strain promotes *in vivo* pathogenesis studies, and importantly, allows for rapid testing of intervention strategies in standard laboratory mice during an expanding pandemic. Utilizing a Venezuelan equine encephalitis virus replicon particle (VRP) system, we vaccinated 10-week-old BALB/c mice against SARS-CoV-2 spike (S), nucleocapsid (N), and GFP as a control,boosted after 3 weeks, and challenged 4 weeks post boost with SARS-CoV-2 MA. Serum samples were taken 3 weeks post boost to measure neutralization titers. Unlike mice vaccinated with GFP or N, serum from S vaccinated mice potently neutralized SARS-CoV-2 reporter virus expressing nanoluciferase (nLUC) (Fig. 4A). Upon challenge with SARS-CoV-2 MA, only those vaccinated with VRP expressing S significantly diminished lung and nasal turbinate titer (Fig. 4B-C).

**Figure 4.**
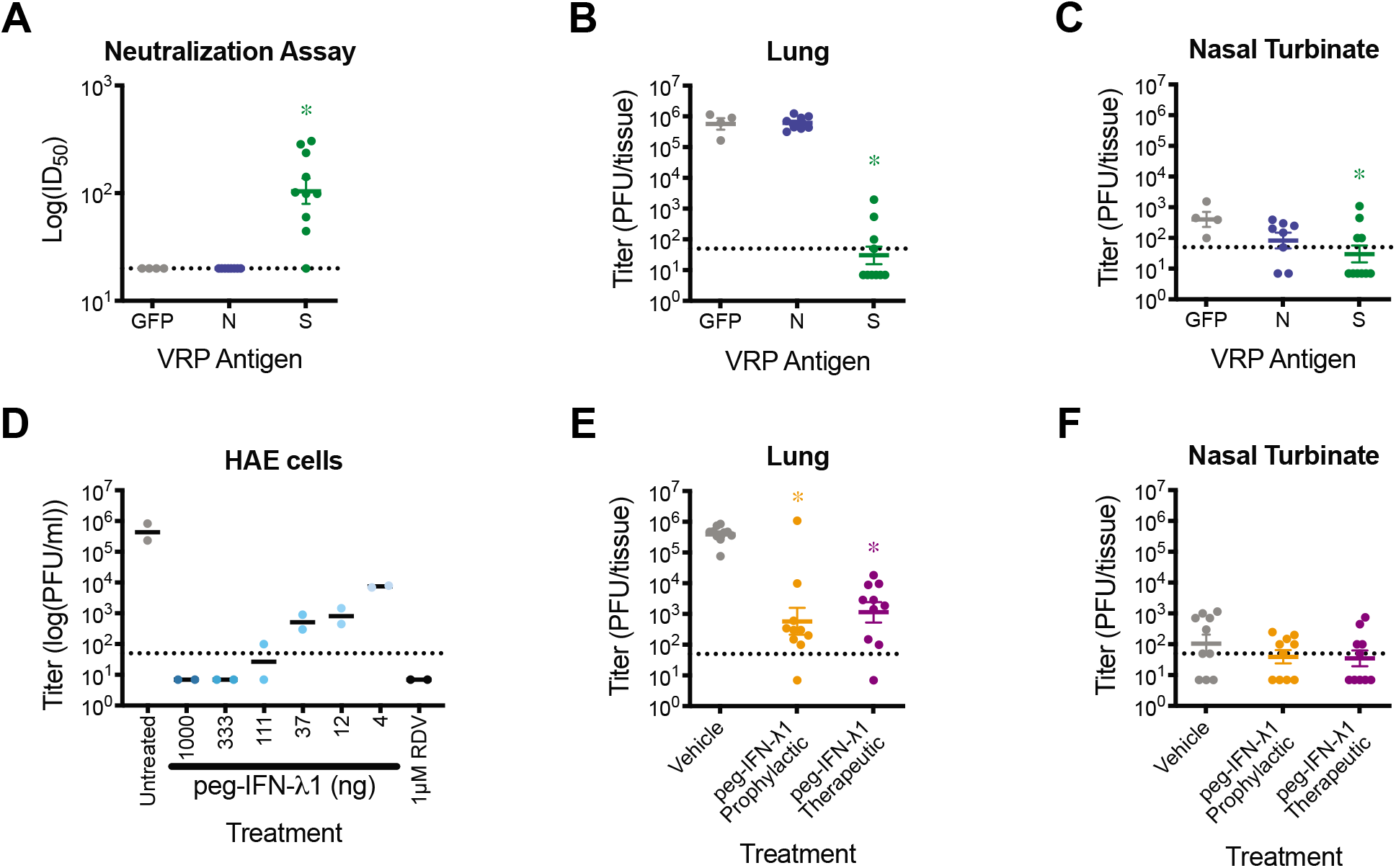
Evaluation of prevention and intervention strategies against SARS-CoV-2 MA infection in mice. **(A-C)** Groups of 10-week-old female BALB/c mice were vaccinated with VRPs expressing wildtype spike (S, green, n=10), nucleocapsid (N, blue, n=8), or GFP (gray, n=4). Mice were boosted 3 weeks after prime immunization, bled 3 weeks post boost for neutralization assays, and challenged with SARS-CoV-2 MA at 4 weeks post boost. **(A)** 50% inhibitory concentration (IC50) values of sera from neutralization of SARS-CoV-2 WT. Dotted line represents limit of detection. Log transformed data analyzed by Kruskal Wallis test followed by Dunnett’s multiple comparisons. **(B)** Lung viral titer. Dotted line represents limit of detection. Undetected samples are plotted at half the limit of detection. Log transformed data analyzed as in **(A)**. **(C)** Nasal turbinate viral titer. Log transformed data analyzed as in **(A)**. **(D)** Human primary airway epithelial cells were pretreated for 24hrs with peg-IFN-λ1 followed by infection with SARS-CoV-2 WT. Infectious virus in apical washes from 48 hours post infection was titered. Remdesivir (RDV) was used as positive control. Dotted line represents limit of detection. Undetected samples are plotted at half the limit of detection. This study was repeated in cells from two unique human donors. **(E-F)** 12-week-old female BALB/c mice were subcutaneously treated with vehicle (gray) or with 2μg peg-IFN-λ1 prophylactically (orange) or therapeutically (purple) and infected with SARS-CoV-2 MA. **(E)** Lung viral titer. Dotted line represents limit of detection. Log transformed data analyzed by Kruskal Wallis test followed by Dunnett’s multiple comparisons. **(F)** Nasal turbinate viral titer. Dotted line represents limit of detection. Log transformed data analyzed by Kruskal Wallis test followed by Dunnett’s multiple comparisons. The line represents the mean and error bars represent standard error of the mean. Asterisk denotes p<0.05.

Interferon lambda is a type III interferon whose receptors are largely limited to epithelial cells, including the lungs, liver, and gastrointestinal tract^16,17^. Treatment with interferons has been employed as pan viral treatment for several viral infections, including trials for the treatment of SARS-CoV-1 and MERS-CoV infections. Pegylated interferon lambda-1 (peg-IFN-λ1) is Phase 3-ready for hepatitis delta virus infection and has been proposed to treat COVID-19 patients^18^. We first sought to determine if peg-IFN-λ1 would initiate an antiviral response capable of inhibiting productive infection of primary human airway epithelial (HAE) cell cultures by SARS-CoV-2. Pretreatment of HAE with peg-IFN-λ1 provided a potent dose dependent reduction in SARS-CoV-2 infectious virus production (Fig 4D). To determine if this *in vitro* antiviral effect would translate to *in vivo* efficacy, we performed prophylactic and therapeutic efficacy studies in BALB/c mice. We subcutaneously administered 2μg peg-IFN-λ1 18hr prior or 12hr after infection with 10^5^ PFU SARS-CoV-2 MA. Both prophylactic and therapeutic administration of peg-IFN-λ1 significantly diminished SARS-CoV-2 MA replication in the lung (Fig. 4E). Peg-IFN-λ1 lowered nasal turbinate titer compared to vehicle treated mice, though not statistically significant due likely due to the limit of detection (Fig. 4F). Altogether, these data demonstrate the utility of this model to rapidly evaluate vaccine and therapeutic drug efficacy in standard laboratory mice. In addition, we show that peg-IFN-λ1 exerts potent antiviral activity against SARS-CoV-2 *in vitro* and can diminish virus replication *in vivo* even when given therapeutically.

## Discussion

Structural studies have solved the interactive networks associated with the SARS-CoV and SARS-CoV-2 RBD interaction residues that bind hACE2^19,20^. Coupling predictive molecular modeling and reverse genetics^21^, we altered the SARS-CoV-2 receptor binding domain (RBD) allowing viral entry via mACE2. Unlike parental wild-type (WT) virus, the resultant recombinant virus, SARS-CoV-2 MA, used mACE2 for infection *in vitro.* SARS-CoV-2 MA replicated in both the upper and lower airways of both 10-week old young adult and 1-year aged BALB/c mice but disease was more severe in aged mice, reproducing the age-related increase in pathogenesis observed in humans. It is remarkable that improved lethal mouse models of SARS-CoV-1 also selected for mutations in key RBD interaction models in *in vivo^22,23^.* Importantly, we show the utility of the SARS-CoV-2 MA model for screening MCMs through vaccine challenge studies and the evaluation of a novel clinical candidate, pegylated interferon lambda-1. Peg-IFN-λ1 is a Phase 3-ready drug in clinical development for hepatitis delta virus infection with a well-established safety and tolerability profile. It has been given to over 3000 patients in the context of 19 clinical trials as a weekly subcutaneous injection, often for 24-48 weeks to treat patients with chronic viral hepatitis^24,25^. For SARS-CoV-2, peg-IFN-λ1 is a promising therapeutic for the treatment of COVID-19 infections^26^, blocks porcine coronavirus replication in gut cells *in vitro^27^*, and SARS-CoV-1 replication in human airway cells^28^. Our data, which demonstrates reduction in SARS-CoV-2 infection in primary human cells and in mice, support further evaluation in primates and in humans. In fact, multiple investigator sponsored studies are underway to evaluate peg-IFN-λ1 for prevention and treatment of SARS-CoV infection (NCT04331899, NCT04343976,NCT04344600, NCT04354259). Thus, we provide a new SARS-CoV-2 animal model that captures multiple aspects of SARS-CoV-2 pathogenesis, providing a high-throughput *in vivo* system to evaluate MCMs and helping fill a critical unmet need for improved human health. In addition, since this model uses standard immune competent laboratory mice, the ease of use, cost and utility is likely greater than that of the current *hACE2* transgenic models. While this model provides a critically necessary tool in COVID-19 countermeasure research, it also provides an important first step in SARS-CoV-2 serial adaptation in mice^23^, potentially selecting for variants that develop more severe pathogenic manifestations of acute respiratory distress syndrome (ARDS), coagulopathy, and other disease outcomes seen in human populations. In addition, the SARS-CoV-2 MA strain can be used to evaluate the role of host innate immune genes and other antiviral defense genes in viral pathogenesis using transgenic and knockout mice.

Although the SARS-CoV-2 MA RBD mutations may attenuate the function of select human monoclonal antibodies or vaccines in mice, this phenotype was not observed with mouse adapted strains of SARS-CoV-1^29-31^. In any event, we demonstrate that our previously described *hACE2* transgenic mouse model supports efficient SARS-CoV-2 replication and pathogenesis *in vivo*, as well^5^. Consequently, this model offers an alternative replication and disease model that uses wildtype SARS-CoV-2, appropriate for evaluating therapeutic antibodies that bind the RBD and other countermeasures. In comparison to SARS-CoV-1 infection, SARS-CoV-2 infection results in a mild bronchiolitis in young mice characterized by efficient virus growth, minimal weight loss and about 40% mortality, the latter phenotype associated with viral invasion of the CNS. In *hACE2* transgenic mice, neuroinvasion is a common pathologic outcome following SARS-CoV infection and a major driver of mortality^5,32^. Together, these data describe critical new animal models,demonstrate countermeasure potential and provide key recombinant viruses for further mouse adaptation, leading to 2^nd^ generation mouse models of human disease.

## Material & Methods

### SARS-CoV-2 receptor binding domain and ACE2 analysis and modelling

Group 2B coronavirus spike and ACE2 amino acid sequences were aligned using Geneious Prime (Version 2020.0.5). Accession numbers used: SARS-CoV-1 Urbani (AY278741), WIV1 (KF367457), SHC014 (KC881005), SARS-CoV-2 (MN985325.1), hACE2 (BAB40370), mACE2 (NP_081562). Protein similarity scores were calculated using BLOSUM62 matrix. Contact residues previously identified by crystal structures^19,20,33^. Structure modelling was performed using Modeller (Version 9.20) and visualized using PyMOL (Version 1.8.6.0).

### Virus and cells

All viruses used were derived from an infectious clone of SARS-CoV-2, which was designed using similar strategies for SARS-CoV and MERS-CoV (14569023, 24043791). The Q498Y/P499T substitutions were generated by site directed mutagenesis using the following primers: Forward: 5’-ATA TGG TTT CTA CAC GAC TAA TGG TGT TGG TTA CCA ACC-3’, Reverse: 5’-TAG TCG TGT AGA AAC CAT ATG ATT GTA AAG GAA AGT AAC AAT TAA AAC CTT C-3’. Viruses were derived following systematic cDNA assembly of the infections clone, followed by *in vitro* transcription and electroporation into Vero E6 cells. Virus stocks were passaged once on Vero E6 cells and titered via plaque assay. Briefly, virus was serial diluted and inoculated onto confluent monolayers of Vero E6 cells, followed by agarose overlay. Plaques were visualized on day 2 post infection via staining with neutral red dye.

Vero E6 cells were maintained in n Dulbecco’s modified Eagle’s medium (DMEM; Gibco), 5% Fetal Clone II serum (FCII, Hyclone), and 1X antibiotic/antimycotic (Gibco). DBT-9 were maintained in DMEM, 10% FCII, and 1X antibiotic/antimycotic.

For single step growth curve, Vero E6 cells were infected at a multiplicity of infection (MOI) of 1 for 1 hour. Inoculum was removed and monolayer was washed twice with PBS, and replace with media. At designated timepoints, media was removed without replacement, and stored at −80°C until titered by plaque assay as described above.

### ACE2 compatibility

For ACE2 receptor usage, non-permissive DBT-9 cells were transfected with pcDNA3.1 empty-vector, pcDNA3.1-hACE2, or pcDNA3.1-mACE2 using lipofectamine 2000 (Invitrogen). 24hrs post transfection, cells were infected at an MOI of 1 for 1 hour. Inoculum was removed and monolayer was washed twice with PBS, and replace with media. At 24hrs post infection, media was removed, and total cellular RNA was collected via TRIzol (Invitrogen) and extracted using Direct-Zol RNA MiniPrep kit (Zymo Research). Viral RNA was quantified via qRT-PCR using TaqMan Fast Virus 1-Step Master Mix (Thermo Fisher Scientific) on a QuantStudio 3 (Applied Biosystems). SARS-CoV-2 RNA was quantified using US Centers of Disease Control and Prevention diagnostic N1 assay: Forward: 5’-GAC CCC AAA ATC AGC GAA AT-3’, probe: 5’-FAM-AC CCC GCA TTA CGT TTG GTG GAC C-BHQ1-3’, reveres: 5’-TCT GGT TAC TGC CAG TTG AAT CTG-3’. Host 18S rRNA was used as housekeeping control (Invitrogen, product number 4319413E). Viral RNA was analyzed using ΔΔCt and fold change over viral RNA in empty-vector transfected cells.

### *In vivo* Infections

hACE2 overexpressing mice were bred and maintained at University of North Carolina at Chapel Hill. BALB/c mice were obtained from Envigo (strain 047). Mice were infected with 10^5^ plaque forming units (PFU) intranasally under ketamine/xylazine anesthesia. Body weight and pulmonary function by whole body plethysmography (Buxco respiratory solutions, DSI Inc.) were monitored daily where indicated. At indicated timepoints, a subset of mice were euthanized by isoflurane overdose and tissue samples were harvested for titer and histopathology analysis. A subset of mice for nasal turbinate histopathology were perfused with 10% phosphate buffered formalin prior to tissue collection. Titer samples were stored at −80°C until homogenized and titered by plaque assay as described above. Histopathology samples were fixed in 10% phosphate buffered formalin for 7 days before paraffin embedding and sectioning. Slide sections were stained with hematoxylin and eosin (H&E) or used for immunohistochemistry for SARS-CoV-2 nucleocapsid.

### Vaccination studies

Non-select BSL2 Venezuelan equine encephalitis virus strain 3526 based replicon particles (VRPs) were generated to express GFP, SARS-CoV-2 spike (S), or nucleocapsid (N) as described previously^34^. Mice were vaccinated via hind footpad infection in 10uL, boosted identically at 3 weeks post prime, and bled via submandibular bleed at 3 weeks to confirm presence of neutralizing antibodies. Neutralizing antibody levels were assessed via neutralization assay using SARS-CoV-2 WT expressing nanoluciferase (nLUC) in place of ORF7a. Briefly, the ORF7a gene of SARS-CoV-2 was removed from the molecular clone and nLUC inserted downstream of the ORF7a transcription regulatory sequence. Recombinant viruses encoding nLUC (SARS-CoV-2 nLUC) were recovered, titered and serial dilutions of sera were incubated with virus for 1 hour at 37°C, then added to monolayers of Vero E6 cells. 48hrs post infection, viral infection was quantified using nLUC activity via Nano-Glo Luciferase Assay System (Promega). 50% inhibitory concentration (IC50) values were calculated from full dilution curves.

Mice were challenged 4 weeks post boost with 10^5^ plaque forming units (PFU) intranasally under ketamine/xylazine anesthesia. Body weight was monitored daily. On day 2 post infection, mice were euthanized by isoflurane overdose and tissue samples were harvested for titer analysis as described above.

### Pegylated-IFN-λ1 treatment *in vitro* and *in vivo*

Peginterferon Lambda-1a was obtained from Eiger BioPharmaceuticals by MTA in GMP prefilled syringes, 0.18 mg/syringe (0.4 mg/mL). Primary HAE cell cultures were obtained from the Tissue Procurement and Cell Culture Core Laboratory in the Marsico Lung Institute/Cystic Fibrosis Research Center at UNC. Human tracheobronchial epithelial cells provided by Dr. Scott Randell were obtained from airway specimens resected from patients undergoing surgery under University of North Carolina Institutional Review Board-approved protocols (#03-1396) by the Cystic Fibrosis Center Tissue Culture Core. Primary cells were expanded to generate passage 1 cells and passage 2 cells were plated at a density of 250,000 cells per well on Transwell-COL (12mm diameter) supports (Corning). Human airway epithelium cultures (HAE) were generated by differentiation at an air-liquid interface for 6 to 8 weeks to form well-differentiated, polarized cultures that resembled *in vivo* pseudostratified mucociliary epithelium (Fulcher et al., 2005). HAEs were treated with a range of peg-IFN-λ1 doses basolaterally for 24hrs prior to infection. 1μM remdesivir was obtained from Gilead Sciences by MTA and was used as a positive control. Cultures were infected at an MOI of 0.5 for 2 hours. Inoculum was removed and culture was washed three times with PBS. At 48hrs post infection, apical washes were taken to measure viral replication via plaque assays as described above. This study was repeated in two separate human donors.

Mice were subcutaneously treated with a single 2μg dose of peg-IFN-λ1 prophylactically at 18hrs prior to infection, therapeutically at 12hrs post infection, or PBS vehicle treated, and infected with 10^5^ plaque forming units (PFU) of SARS-CoV-2 MA intranasally under ketamine/xylazine anesthesia. Body weight was monitored daily. On day 2 post infection, mice were euthanized by isoflurane overdose and tissue samples were harvested for titer analysis as described above.

### Histopathology and antigen staining

Lungs were fixed for 7 days in 10% phosphate buffered formalin, paraffin embedded, and sectioned at 4μm. Serial sections were stained with hematoxylin and eosin, and stained for immunohistochemistry for SARS-CoV-2 nucleocapsid using a monoclonal anti-SARS-CoV-1 nucleocapsid antibody (NB100-56576, Novus Biologicals) on deparaffinized sections on the Ventana Discovery Ultra platform (Roche). Photomicrographs were captured on an Olympus BX43 light microscope at 200X magnification with a DP27 camera using cellSens Dimension software.

### Data analysis and presentation

All data visualize and analyzed in Prism (version 8.4.2). Non-parametric tests were performed as described in figure legends. Figures arranged in Adobe Illustrator (version 24.1).

### Ethics and containment procedures

All recombinant viruses were approved by the University of North Carolina at Chapel Hill Institutional Review Board under Schedule G 73790. All animal work was approved by Institutional Animal Care and Use Committee at University of North Carolina at Chapel Hill under protocol 19-168 according to guidelines outlined by the Association for the Assessment and Accreditation of Laboratory Animal Care and the U.S. Department of Agriculture. All virus studies were performed in animal biosafety level 3 facilities at University of North Carolina at Chapel Hill.

## Acknowledgements

This project was funded in part by the National Institute of Allergy and Infectious Diseases, National Institutes of Health, Department of Health and Human Service award: 1U19 AI142759 (Antiviral Drug Discovery and Development Center awarded to R.S.B); 5R01AI132178 (partnership grant awarded to T.P.S. and R.S.B) and an animal models contract from the NIH (HHSN272201700036I). D.R.M is funded by an NIH NIAID T32 AI007151 and a Burroughs Wellcome Fund Postdoctoral Enrichment Program Award. The Marsico Lung Institute Tissue Procurement and Cell Culture Core is supported by NIH grant DK065988 and Cystic Fibrosis Foundation grant BOUCHE15RO. Animal histopathology service was performed by Dawud Hilliard and Ling Wang in the Animal Histopathology & Laboratory Medicine Core at the University of North Carolina, which is supported in part by an NCI Center Core Support Grant (5P30CA016086-41) to the UNC Lineberger Comprehensive Cancer Center.

## Author contribution

K.H.D III and S.R.L. designed and conducted *in vitro* and animal experiments, analyzed data, generated figures, and wrote manuscript. A.S. conducted animal experiments, C.E.E. generated VRP vaccine and vaccinated mice. D.R.M. conducted neutralization assay and analyzed data. S.A.M imaged and analyzed histology. A.W., K.E.G., and A.J.B. assisted with animal experiments. B.L.Y. and Y.J.H. performed cloning. L.E.J.A and E.H. assisted with transfection experiments. M.D.B., I.C.C., and J.S.G. provided resources and helped design and analyze experiments with peg-IFN-λ1. L.E.G conducted experiments, analyzed data, and edited manuscript. T.P.S. conducted experiments, analyzed data, and wrote manuscript. R.S.B supervised the project and wrote the manuscript.

---

**Supplemental Figure 1:**
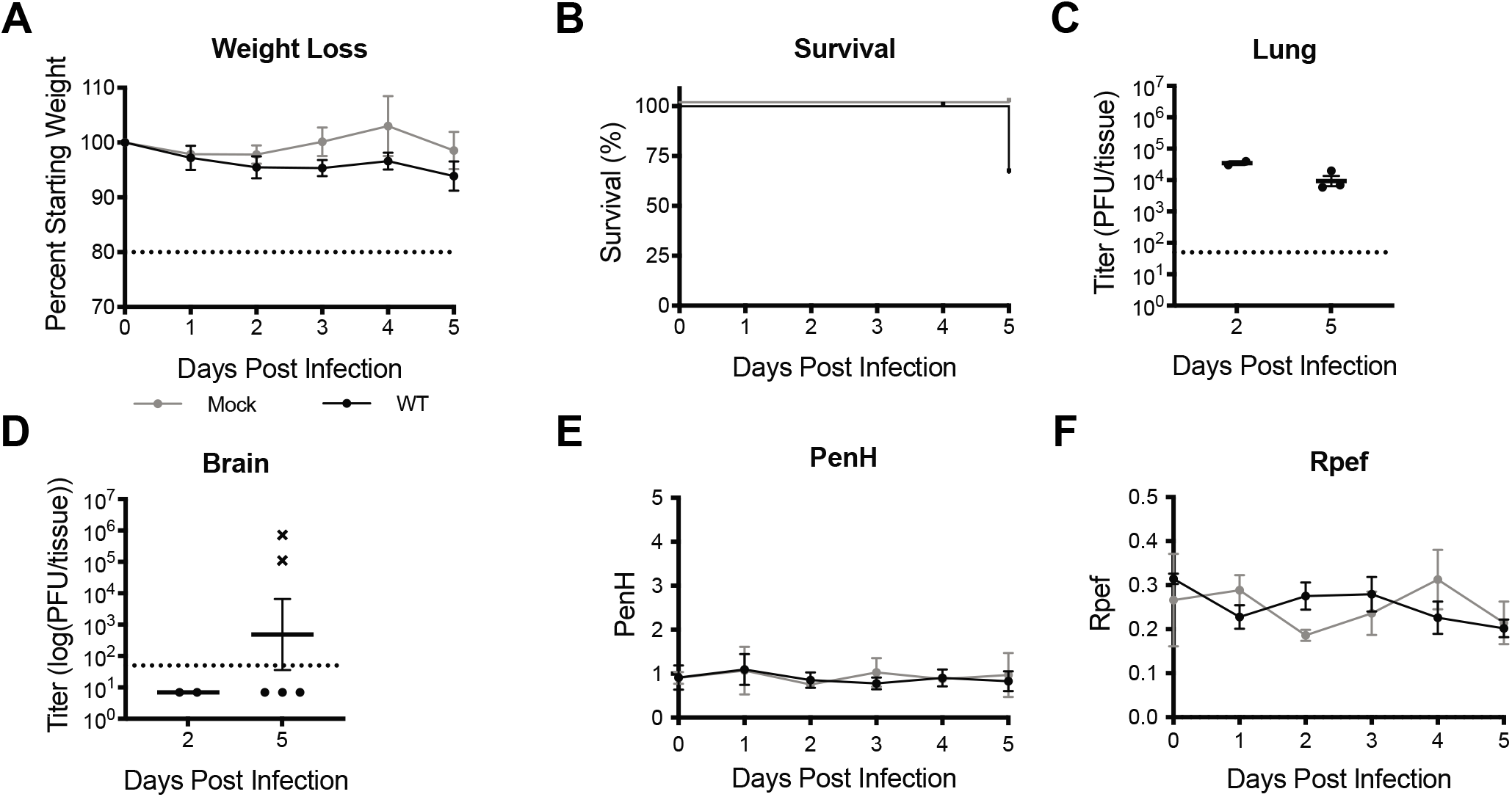
SARS-CoV-2 infection in *hACE2* transgenic mice. **(A)** Percent starting weight. Percent starting weight. Dotted line represents weight loss criteria for humane euthanasia. Data analyzed by mixed effects analysis followed by Sidak’s multiple comparisons. **(B)** Survival. **(C)** Lung viral titer. **(D)** Brain viral titer. Dotted line represents limit of detection. Undetected samples are plotted at half the limit of detection. ‘x’ symbol indicates mice that succumbed to infection. **(E-F)** Whole body plethysmography assessing pulmonary function for PenH **(E)** and Rpef **(F)**. Data analyzed by mixed effects analysis followed by Sidak’s multiple comparisons. The line represents the mean and error bars represent standard error of the mean. Asterisk denotes p<0.05.

**Supplemental Figure 2:**
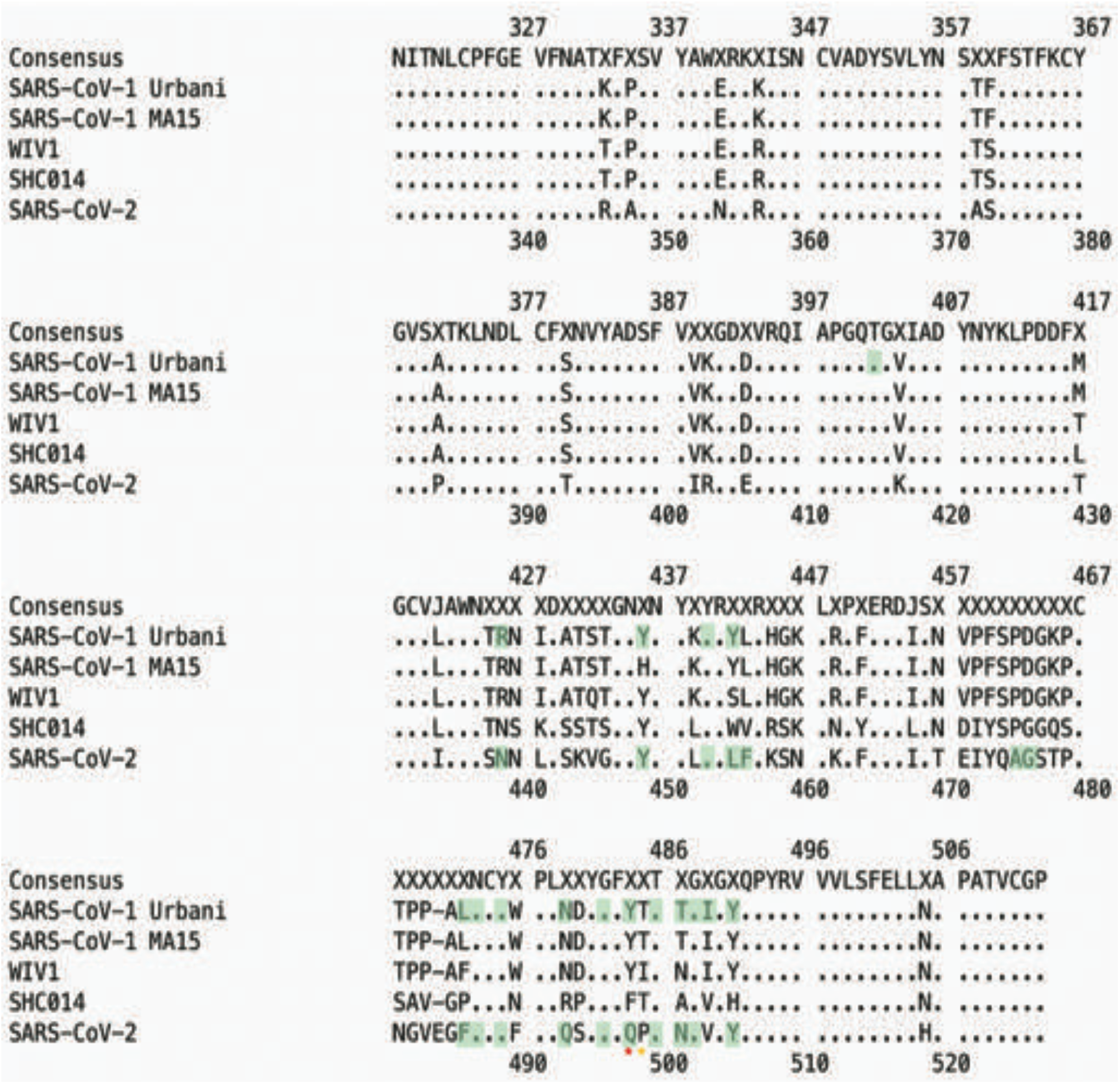
Group 2B coronavirus spike receptor binding domain alignment. Amino acid positions are numbered above in reference to SARS-CoV-1, and below in reference to SARS-CoV-2. Green highlighted residues are hACE2 contacts as determined by published crystal structures.

**Supplemental Figure 3:**
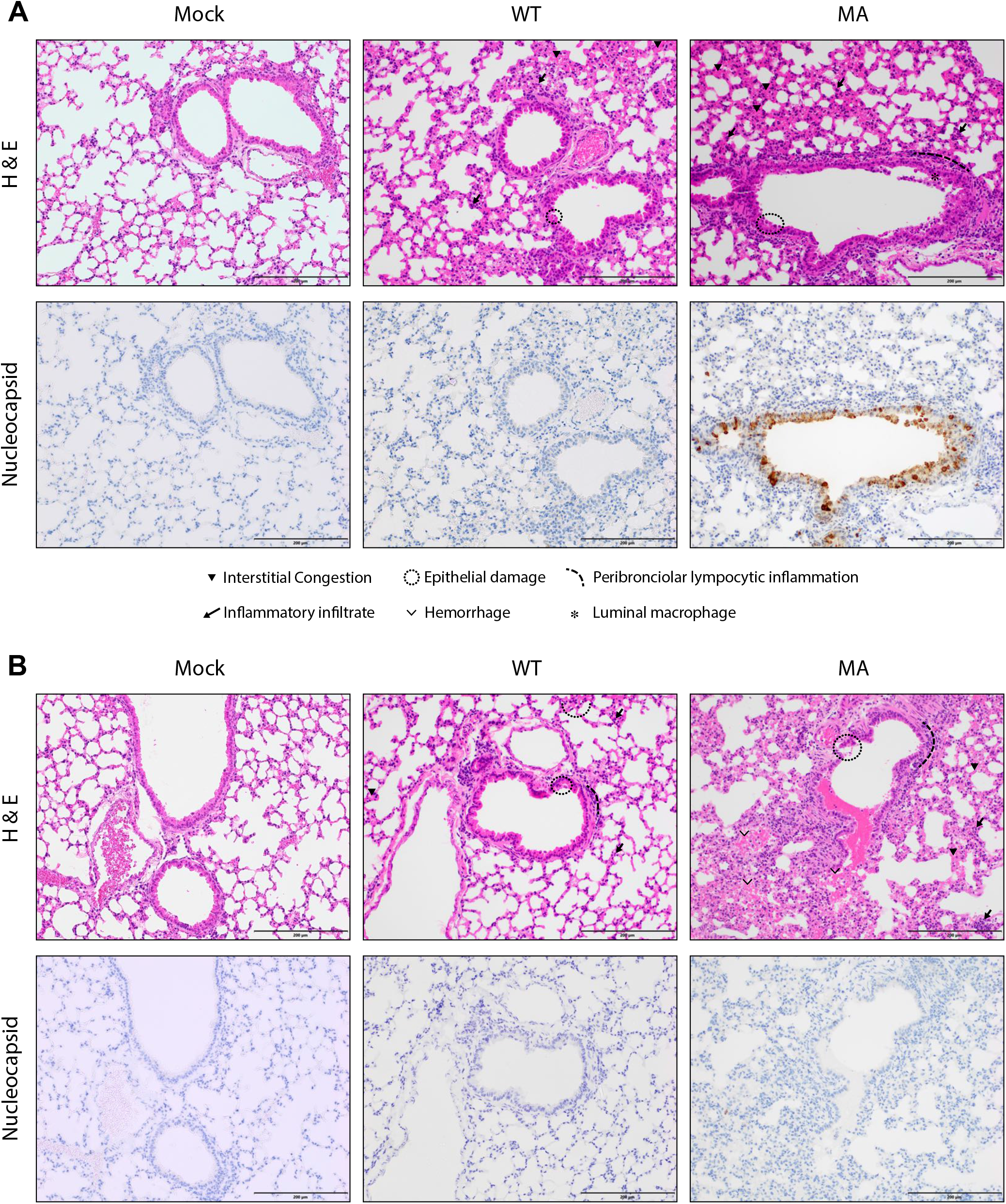
Histology and SARS-CoV-2 MA antigen staining in young BALB/c lungs. Representative images of lung sections from 12-week-old BALB/c mice from Fig. 2. (A) 2dpi. (B) 4dpi. Top panels are hematoxylin and eosin (H & E). Bottom panels are immunohistochemistry staining for SARS-CoV-2 nucleocapsid protein, counterstained with hematoxylin.

**Supplemental Figure 4:**
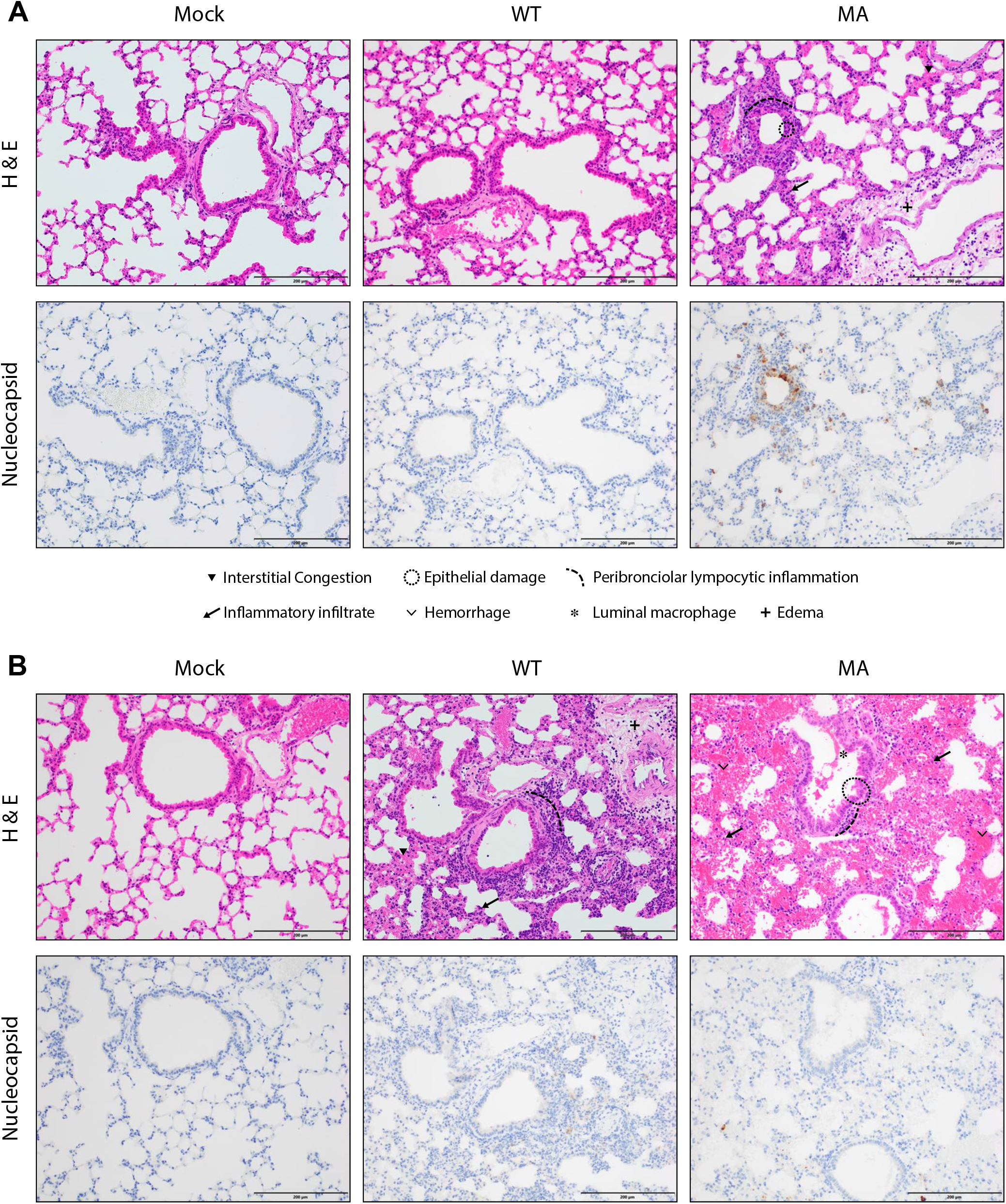
Histology and SARS-CoV-2 MA antigen staining in old BALB/c lungs. Representative images of lung sections from 1-year-old BALB/c mice from Fig. 2. (A) 2dpi. (B) 4dpi. Top panels are hematoxylin and eosin (H & E). Bottom panels are immunohistochemistry staining for SARS-CoV-2 nucleocapsid protein, counterstained with hematoxylin.

## References

1 Sheahan, T. P. et al. Broad-spectrum antiviral GS-5734 inhibits both epidemic and zoonotic coronaviruses. Sci Transl Med 9, doi:10.1126/scitranslmed.aal3653 (2017).

2 Sheahan, T. P. et al. An orally bioavailable broad-spectrum antiviral inhibits SARS-CoV-2 in human airway epithelial cell cultures and multiple coronaviruses in mice. Sci Transl Med 12, doi:10.1126/scitranslmed.abb5883 (2020).

3 Sheahan, T. P. et al. Comparative therapeutic efficacy of remdesivir and combination lopinavir, ritonavir, and interferon beta against MERS-CoV. Nat Commun 11, 222, doi:10.1038/s41467-019-13940-6 (2020).

4 Zhou, P. et al. A pneumonia outbreak associated with a new coronavirus of probable bat origin. Nature 579, 270–273, doi:10.1038/s41586-020-2012-7 (2020).

5 Menachery, V. D. et al. SARS-like WIV1-CoV poised for human emergence. Proc Natl Acad Sci U S A 113, 3048–3053, doi:10.1073/pnas.1517719113 (2016).

6 Menachery, V. D., Gralinski, L. E., Baric, R. S. & Ferris, M. T. New Metrics for Evaluating Viral Respiratory Pathogenesis. PLoS One 10, e0131451, doi:10.1371/journal.pone.0131451 (2015).

7 Cockrell, A. S. et al. A mouse model for MERS coronavirus-induced acute respiratory distress syndrome. Nat Microbiol 2, 16226, doi:10.1038/nmicrobiol.2016.226 (2016).

8 Wang, D. et al. Clinical Characteristics of 138 Hospitalized Patients With 2019 Novel Coronavirus-Infected Pneumonia in Wuhan, China. JAMA, doi:10.1001/jama.2020.1585 (2020).

9 de Wit, E., van Doremalen, N., Falzarano, D. & Munster, V. J. SARS and MERS: recent insights into emerging coronaviruses. Nat Rev Microbiol 14, 523–534, doi:10.1038/nrmicro.2016.81 (2016).

10 Roberts, A. et al. Aged BALB/c mice as a model for increased severity of severe acute respiratory syndrome in elderly humans. J Virol 79, 5833–5838, doi:10.1128/JVI.79.9.5833-5838.2005 (2005).

11 Sheahan, T. et al. Successful vaccination strategies that protect aged mice from lethal challenge from influenza virus and heterologous severe acute respiratory syndrome coronavirus. J Virol 85, 217–230, doi:10.1128/JVI.01805-10 (2011).

12 Bolles, M. et al. A double-inactivated severe acute respiratory syndrome coronavirus vaccine provides incomplete protection in mice and induces increased eosinophilic proinflammatory pulmonary response upon challenge. J Virol 85, 12201–12215, doi:10.1128/JVI.06048-11 (2011).

13 Sheahan, T. et al. MyD88 is required for protection from lethal infection with a mouse-adapted SARS-CoV. PLoS Pathog 4, e1000240, doi:10.1371/journal.ppat.1000240 (2008).

14 Frieman, M. B. et al. SARS-CoV pathogenesis is regulated by a STAT1 dependent but a type I, II and III interferon receptor independent mechanism. PLoS Pathog 6, e1000849, doi:10.1371/journal.ppat.1000849 (2010).

15 Gralinski, L. E. et al. Complement Activation Contributes to Severe Acute Respiratory Syndrome Coronavirus Pathogenesis. mBio 9, doi:10.1128/mBio.01753-18 (2018).

16 Kotenko, S. V. et al. IFN-lambdas mediate antiviral protection through a distinct class II cytokine receptor complex. Nat Immunol 4, 69–77, doi:10.1038/ni875 (2003).

17 Sheppard, P. et al. IL-28, IL-29 and their class II cytokine receptor IL-28R. Nat Immunol 4, 63–68, doi:10.1038/ni873 (2003).

18 Elazar, M. & Glenn, J. S. Emerging concepts for the treatment of hepatitis delta. Curr Opin Virol 24, 55–59, doi:10.1016/j.coviro.2017.04.004 (2017).

19 Lan, J. et al. Structure of the SARS-CoV-2 spike receptor-binding domain bound to the ACE2 receptor. Nature, doi:10.1038/s41586-020-2180-5 (2020).

20 Shang, J. et al. Structural basis of receptor recognition by SARS-CoV-2. Nature, doi:10.1038/s41586-020-2179-y (2020).

21 Schabort, I., Odendaal, H. J., Lombard, C. J. & Bredell, L. Comparison between umbilical artery and vein endogenous digoxin-like immuno-active factor levels in normal and preeclamptic patients. S Afr Med J 79, 197–199 (1991).

22 Frieman, M. et al. Molecular determinants of severe acute respiratory syndrome coronavirus pathogenesis and virulence in young and aged mouse models of human disease. J Virol 86, 884–897, doi:10.1128/JVI.05957-11 (2012).

23 Roberts, A. et al. A mouse-adapted SARS-coronavirus causes disease and mortality in BALB/c mice. PLoS Pathog 3, e5, doi:10.1371/journal.ppat.0030005 (2007).

24 Muir, A. J. et al. A randomized phase 2b study of peginterferon lambda-1a for the treatment of chronic HCV infection. J Hepatol 61, 1238–1246, doi:10.1016/j.jhep.2014.07.022 (2014).

25 Chan, H. L. Y. et al. Peginterferon lambda for the treatment of HBeAg-positive chronic hepatitis B: A randomized phase 2b study (LIRA-B). J Hepatol 64, 1011–1019, doi:10.1016/j.jhep.2015.12.018 (2016).

26 Andreakos, E. & Tsiodras, S. COVID-19: lambda interferon against viral load and hyperinflammation. EMBO Mol Med, doi:10.15252/emmm.202012465 (2020).

27 Li, L. et al. IFN-lambda preferably inhibits PEDV infection of porcine intestinal epithelial cells compared with IFN-alpha. Antiviral Res 140, 76–82, doi:10.1016/j.antiviral.2017.01.012 (2017).

28 Mordstein, M. et al. Lambda interferon renders epithelial cells of the respiratory and gastrointestinal tracts resistant to viral infections. J Virol 84, 5670–5677, doi:10.1128/JVI.00272-10 (2010).

29 Rockx, B. et al. Escape from human monoclonal antibody neutralization affects in vitro and in vivo fitness of severe acute respiratory syndrome coronavirus. J Infect Dis 201, 946–955, doi:10.1086/651022 (2010).

30 Sui, J. et al. Effects of human anti-spike protein receptor binding domain antibodies on severe acute respiratory syndrome coronavirus neutralization escape and fitness. J Virol 88, 13769–13780, doi:10.1128/JVI.02232-14 (2014).

31 Rockx, B. et al. Structural basis for potent cross-neutralizing human monoclonal antibody protection against lethal human and zoonotic severe acute respiratory syndrome coronavirus challenge. J Virol 82, 3220–3235, doi:10.1128/JVI.02377-07 (2008).

32 McCray, P. B., Jr. et al. Lethal infection of K18-hACE2 mice infected with severe acute respiratory syndrome coronavirus. J Virol 81, 813–821, doi:10.1128/JVI.02012-06 (2007).

33 Li, F., Li, W., Farzan, M. & Harrison, S. C. Structure of SARS coronavirus spike receptor-binding domain complexed with receptor. Science 309, 1864–1868, doi:10.1126/science.1116480 (2005).

34 Agnihothram, S. et al. Development of a Broadly Accessible Venezuelan Equine Encephalitis Virus Replicon Particle Vaccine Platform. J Virol 92, doi:10.1128/JVI.00027-18 (2018).

